# De novo birth of functional, human-specific microproteins

**DOI:** 10.1101/2021.10.01.462744

**Authors:** Nikolaos Vakirlis, Kate M. Duggan, Aoife McLysaght

## Abstract

We now have a growing understanding that functional short proteins can be translated out of small Open Reading Frames (sORF). Such “microproteins” can perform crucial biological tasks and can have considerable phenotypic consequences. However, their size makes them less amenable to genomic analysis, and their evolutionary origins and conservation are poorly understood. Given their short length it is plausible that some of these functional microproteins have recently originated entirely de novo from non-coding sequence. Here we test the possibility that de novo gene birth can produce microproteins that are functional “out-of-the-box”. We reconstructed the evolutionary origins of human microproteins previously found to have measurable, statistically significant fitness effects. By tracing the appearance of each ORF and its transcriptional activation, we were able to show that, indeed, novel small proteins with significant phenotypic effects have emerged de novo throughout animal evolution, including many after the human-chimpanzee split. We show that traditional methods for assessing the coding potential of such sequences often fall short, due to the high variability present in the alignments and the absence of telltale evolutionary signatures that are not yet measurable. Thus we provide evidence that the functional potential intrinsic to sORFs can be rapidly, and frequently realised through de novo gene birth.

## Introduction

It is now a well-established biological fact that many more ORFs are translated than those traditionally annotated as protein-coding^1^. Most of these so called “noncanonical” ORFs, such as ones found on long noncoding RNAs, are small, typically <300 nucleotides. While most of these are plausibly just biological noise, many encode functional microproteins^2^. Microproteins perform diverse functions through various mechanisms: some, encoded by upstream ORFs (uORFs), exert translational control over the main ORF of the transcript^3^; while others interact with larger protein complexes or with cellular membranes^4,5^. Microproteins have long been overlooked in genomic studies, mostly due to technical limitations linked to their small size^6^. But there is now increasing interest and investment towards identifying them and understanding their functions and possible roles in health and disease^7,8^. Well studied examples of functional human microproteins include NoBody^9^, PIGBOS^10^ and Myoregulin^11^, while many more have been identified in other species such as mouse^12^, plants^13^, bacteria^14^ and elsewhere^15^. Microproteins have been observed to be highly conserved over long evolutionary times in animals and in plants^16–18^, but they can also be evolutionarily novel^5,19^.

Evolutionarily novel genes can evolve out of preexisting ones through sequence divergence^20–22^ (preceded by duplication or not), but they can also emerge entirely de novo, out of ancestrally non-genic genomic regions^23^. The process of de novo gene birth, as the latter is called, has now been studied extensively in multiple species such as yeast^24–26^, mouse^27^, flies^28^, fish^29,30^, rice^31^ nematodes^32^ and human^33^. In human, early studies relying on gene annotations^33–36^ established that de novo genes can indeed form, even in as short an evolutionary timeframe as the split of human from chimpanzee. Later studies adopted broader search strategies, starting from entire transcriptomes and incorporating ribosome profiling data to identify translated ORFs^37,38^.

While many studies have addressed the conservation of human microproteins, their modes of origin, de novo or otherwise, have not been systematically investigated. Indeed, conservation is widely used as a coding/functional signature and hence non-conserved, novel ORFs are excluded from most studies. However, it is plausible and even intuitive that novel genes may first arise as ORFs coding for very small proteins. Given the fact that de novo gene birth seems to consistently result in short ORF sequences^23,39^ (at least initially), and that microproteins perform functions out of simple structures, it follows that human microproteins could have recently emerged de novo and already assumed selectively relevant cellular functions. Thus, the study of microproteins and of de novo emerged genes naturally intersect.

Here, we leveraged the depth of a recently published dataset of human microproteins translated from non-canonical ORFs^40^ to look for such evolutionary birth events. More specifically, we sought to understand whether de novo gene birth can produce proteins that are functional “out-of-the-box”, i.e. immediately after they come into existence. Answering this question is doubly important: for our understanding of the intriguing, and still largely mysterious phenomenon of de novo gene birth, but also for our appreciation of the full functional potential of the human genome.

## Results

### The reconstructed evolutionary origins of human microproteins

A recent rigorous analysis of ribosome profiling data by Chen et al. enabled ORF translation to be inferred with high-confidence for hundreds of human non-canonical ORFs^40^. We used these data and focused on ORFs that did not overlap canonical, coding ones: i.e., are either located on transcripts previously annotated as non-coding (‘new’); or upstream/downstream of known coding ORFs (‘upstream’ and ‘downstream’ respectively); or new mRNA isoforms of an annotated protein-coding gene, but where the new isoform lacked an annotated coding ORF (‘new_iso’), labeled as per classification of Chen et al. (see Methods). Furthermore, we only kept those ORFs which we could unambiguously match to ORFs identified and analysed by Hon et al.^41^ in their comprehensive human transcriptome atlas with accurate 5’ ends (FANTOM CAT). A total of 715 ORFs were included in the final dataset (499 “upstream”, 179 “new”, 32 “new_iso” and 5 “downstream”). They range from 33nt to 3,825nt in length, with a median of 81nt.

For each ORF, we searched for its orthologous chromosomal region in the genomes of 99 other vertebrate species (see Methods, Supp. Table 1). For each ORF, the orthologous nucleotide sequences were aligned, and we constructed a phylogenetic tree for each, following the species tree topology, to estimate the branch lengths. Finally Ancestral Sequence Reconstruction (ASR) was performed, and the presence or not of an ORF at each human ancestral node was inferred. To decide whether an ORF was present or not, we applied a length ratio cut-off (length of ancestral ORF/length of full ancestral sequence) of 70% (see Methods). Whenever an ancestor lacking an intact ORF was found to precede all ancestors with intact ORFs, it was deemed as de novo ORF origination (Figure 1A).

**Figure 1.**
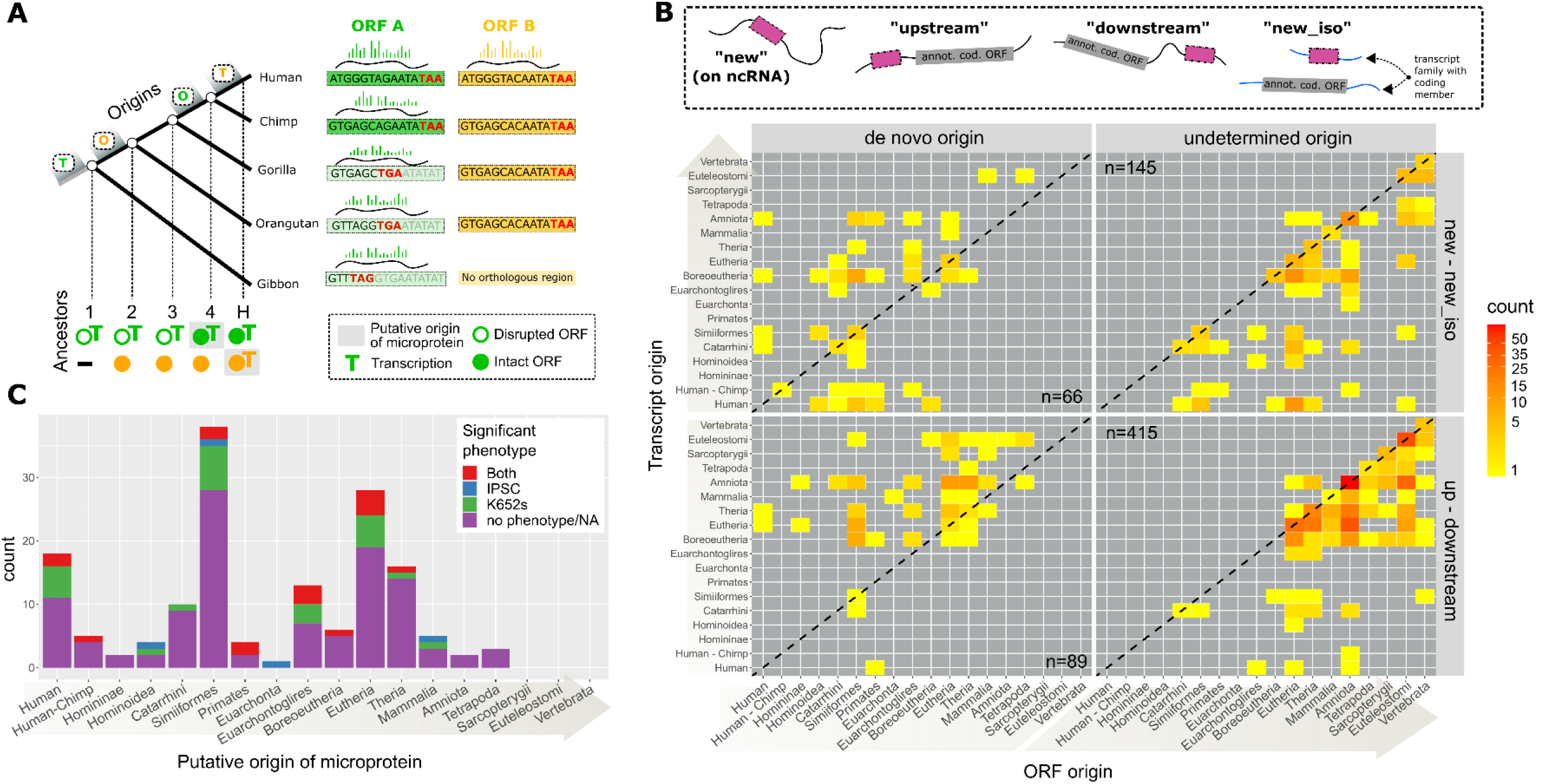
**A**: Graphical example of reconstruction of the timings of origin of the ORF and of transcription, for two hypothetical human microproteins. Human ORF A, is intact in Chimp, but disrupted by a premature stop codon in the other species. Since the orthologous genomic region has been identified in all four species, we can reconstruct the sequence of all 4 ancestors, and determine whether the ORF is intact or not. In this case, the ORF is not intact (disrupted) in ancestors 1,2,3 and intact in ancestor 4. We can thus infer that the ORF emerged de novo and place the node of origin of the ORF (green “O”) between ancestors 3 and 4 (for practical purposes we use the age of ancestor 4). The region orthologous to ORF A has been found to be transcribed in all 4 species, hence the node of origin of transcription is placed at the earliest ancestor (green “T”). The putative origin of the microprotein (grey rectangle) is then calculated as the most recent of the two, which is ancestor 4. ORF B on the other hand, is intact in all 4 species where the orthologous region can be identified. ASR estimates that the ORF was intact in ancestors 2,3,4, but no ancestor prior to that one can be inferred. Hence this is a case of “undetermined origin”. No transcript has been found in any of the other species, thus transcription is inferred to be human-specific. The putative origin for ORF B is therefore the human branch. **B**: Distribution of the phylogenetic origins of ORFs and transcripts in the two broad categories of ORFs. Species and age corresponding to each node can be found in Supp. Figure 2. Nodes are ordered from recent to ancient. **C**: Numbers of de novo originated microproteins with and without significant phenotype in the two cell lines as estimated by Chen et al, grouped by their inferred putative origin. 20 de novo ORFs that have no associated phenotype data are included in the “no phenotype” class.

In total, de novo origin was inferred for 155 ORFs. To assess the influence of the length ratio parameter, we tested alternative values: one where de novo attribution was stricter (50%); and one more relaxed (80%). The same node of origin was inferred for 102/155 and 148/155 de novo ORFs, using the stricter and the more relaxed cut-off respectively (the differences in inferred ages of origin for de novo ORFs can be found in Supp. Figure 1). Thus, approximately 2/3 of our de novo inferred ORFs are entirely robust to this parameter, and for an additional 14 of them the alternative parameter only changed the date by one or two nodes of the tree.

The presence or absence of transcription in the orthologous region of 92/99 vertebrate species was inferred by examining the overlap with annotated transcripts and the similarity to known transcript sequences (see Methods). Transcription was inferred to have originated at the most recent common ancestor of all species in which transcription of the region was detected. Transcription origin, ORF origin as well as all other data gathered for the 715 ORFs can be found in Supp. Table 2.

In Figure 1B we have plotted the distribution of ORF and transcript origination nodes on the vertebrate phylogeny. The origin of the ORF in most cases in the “undetermined origin” category is biased towards the oldest nodes. This result is expected and most likely reflects the limits of homology detection over time due to sequence divergence^20^. The fact that an ORF-first origin (datapoints below the diagonal) is more prevalent for these is probably also an artefact due to our limited capacity to identify distant homologous transcripts. On the other hand, for those cases for which a de novo origin can be inferred, we see a prevalence of RNA-first origin for those in the “up – downstream” group. This is to be expected as these ORFs have mostly formed on pre-existing mRNAs. For those in the “new - new_iso” group, the situation appears more balanced, with a mix of RNA-first and ORF-first cases. We observe a small number of exceptional cases where an ORF has been sufficiently conserved to allow homology detection since as far back as the split of the Boreoeutheria, but we only see evidence for transcription in the human branch. Given that transcript discovery and annotation is far from complete outside of model organisms, it is possible that some of these cases could indeed be false positives, explained by a lack of identified transcripts in species other than human. Alternatively, some could correspond to spurious ORFs that simply happen to overlap with hyper conserved elements such as enhancers.

We used the data on ORF conservation and detectable transcription to infer the branch of origin of the 155 de novo origin microproteins. In order for the microprotein to be produced, there must be both an ORF and transcription, so we define the origin of the microprotein as the earliest node where both are detected (shaded boxes in Figure 1A, henceforth the term “putative origin” will be used for this).

### Evidence for the biological significance of de novo emerged microproteins

The functional relevance of young de novo originated ORFs is debated. We thus asked whether any of our recently de novo emerged, robustly translated microproteins was found to be functional. For 44/155 de novo originated microproteins, CRISPR-Cas based disruption of their ORF was found to have statistically significant fitness effects in two cell lines (iPSC and K562) according to the strict criteria of Chen et al. This proportion is statistically indistinguishable from that for microproteins of undetermined origin (156/560, X^2^ P-value=0.98). The putative origin and knockout phenotype for each of the 155 de novo emerged microproteins can be found in Figure 1C.

Our results suggest that there has been ongoing de novo birth of functional microproteins since the early evolution of mammals. At least one such microprotein has originated at 12/13 nodes going back to the mammalian ancestor. The absence in older nodes can be explained by the overall low number of de novo genes identified there, which in turn is due to the long evolutionary times. There are 14 de novo emerged, functional microproteins that come from ORFs found on lncRNAs and have a putative origin within the past 43my (since the ancestor of higher primates, Simiiformes). Notably, 6 of these functional microproteins (4 if we remove overlapping ones, on 4 distinct transcripts), have a putative origin after the split of human and chimpanzee. All 6 of these cases are ORF-first, with the most recent one having an estimated ORF origin at the Hominoidea. This observation provides strong support for the hypothesis that de novo emerged microproteins have a ready route to biological significance and may indeed be functional “out of the box”.

Numbers of de novo microproteins with significant phenotypes across ages correlate strongly with those without a phenotype (Spearman’s Rho=0.65, P-value=0.0092, excluding the nodes Sarcopterygii, Euteleostomi and Vertebrata, for which no de novo ORFs were found). This suggests that a steady proportion of de novo microproteins acquire biological significance. Additionally, it implies that a more powerful search for novel microproteins may uncover further functional examples, and that the observed numbers simply reflect the limited sampling of cell types and growth conditions tested.

The fact that some of the microproteins with measurable phenotypes have recently emerged de novo and are entirely novel, further reinforces the fact that evolutionary conservation and coding signals alone do not reveal the full repertoire of protein-coding genes in a genome. Indeed, out of the 44 de novo emerged microproteins with functional evidence, none were predicted as coding by PhyloCSF^42^, a widely used comparative genomics tool that determines the likelihood that a sequence is protein-coding based on a nucleotide multiple sequence alignment (see refs ^40,41^ and Supp. Table 2). None were predicted as coding by RNAcode^43^ either (ref ^41^ and Supp. Table 2) and only 4/44 were predicted to be coding based on the ribosome profiling measure (FLOSS score)^41^. Only 2 have a CPAT^44^ coding probability higher than 0.5 when calculated over the ORF sequence only (mean of 0.093) and only 4 are predicted by CPAT to be coding based on analysis of the entire transcript (calculated by Hon et al. see Supp. Table 2).

So, are the fitness effects observed really due to the absence of the expressed protein, or could they be coming from the regulatory or RNA level? Chen et al. performed rescue experiments for 9 “upstream” and 7 “new” ORFs where the ORF peptide was ectopically expressed, as well as controls in which the start codon of the expressed ORF was removed. In all cases, the growth phenotype was rescued, and it was shown that the rescue was dependent on the presence of the start codon. Out of the 7 validated “new” ORFs, 5 are included in our analysis. Only two, with putative origin at Tetrapoda and Amniota show signs of being coding (CPAT and PhyloCSF), while the other three are all much more recent with putative origin at Hominoidea (de novo), Eutheria (undetermined) and human (undetermined). Similar results are found for “upstream” ORFs. Out of the 7 we analysed, 5 show coding signatures and they are all at least as ancient as mammals. The only young one, CATP00000415540.1, with a de novo origin at the Simiiformes, entirely lacks coding signatures. While more validation experiments will be necessary, these results seem to confirm that the fitness effects of these young, not characteristically coding ORFs, are indeed linked to the action of the protein.

Comparative methods such as PhyloCSF should be applied with caution. One difficulty in employing and interpreting PhyloCSF scores in cases such as ours is that failure to recognize the recent de novo origin of genes may result in the inclusion of sequences from lineages that diverged before the gene origin and where the ORF is not present. This can negatively bias the coding assessment when the algorithm (correctly) infers a lack of coding potential in a large number of the provided sequences. Frameshifts, which should be more common in evolutionarily recent, less constrained coding sequences, can further complicate coding prediction.

Under this rationale, we hypothesized that considering the phylogenetic origin of each ORF and the conservation of the reading-frame in each alignment might ameliorate coding signature detection. We thus applied PhyloCSF in codon-aware alignments comprising only species descending from the predicted node of origin of the ORF (origin of transcription is not taken into account for this specific calculation, see Methods). We then counted the number of ORFs predicted to be coding by each study, taking a frequently used cut-off of PhyloCSF score ≥ 41^41,45^. We obtained 62 coding ORFs, that is, 2.8 times more than Chen et al. (22) and 1.8 times more than Hon et al. (35). Half (31/62) are not predicted as coding by either previous study. Importantly, these 31 ones unique to this study, are biased with respect to significant phenotype (19/31, X^2^ test, P-value = 0.00015). These results, grouped in four broad classes of age of putative origin, are shown in Figure 2A. A similar difference in coding predictions is observed when relaxing the score cut-off to ≥10 (75 vs. 42 and 48). Finally, our approach is the only one that identifies any de novo originated microprotein with phenotypic effects as coding (4 vs 0 and 0), arguably the toughest and most critical cases.

**Figure 2:**
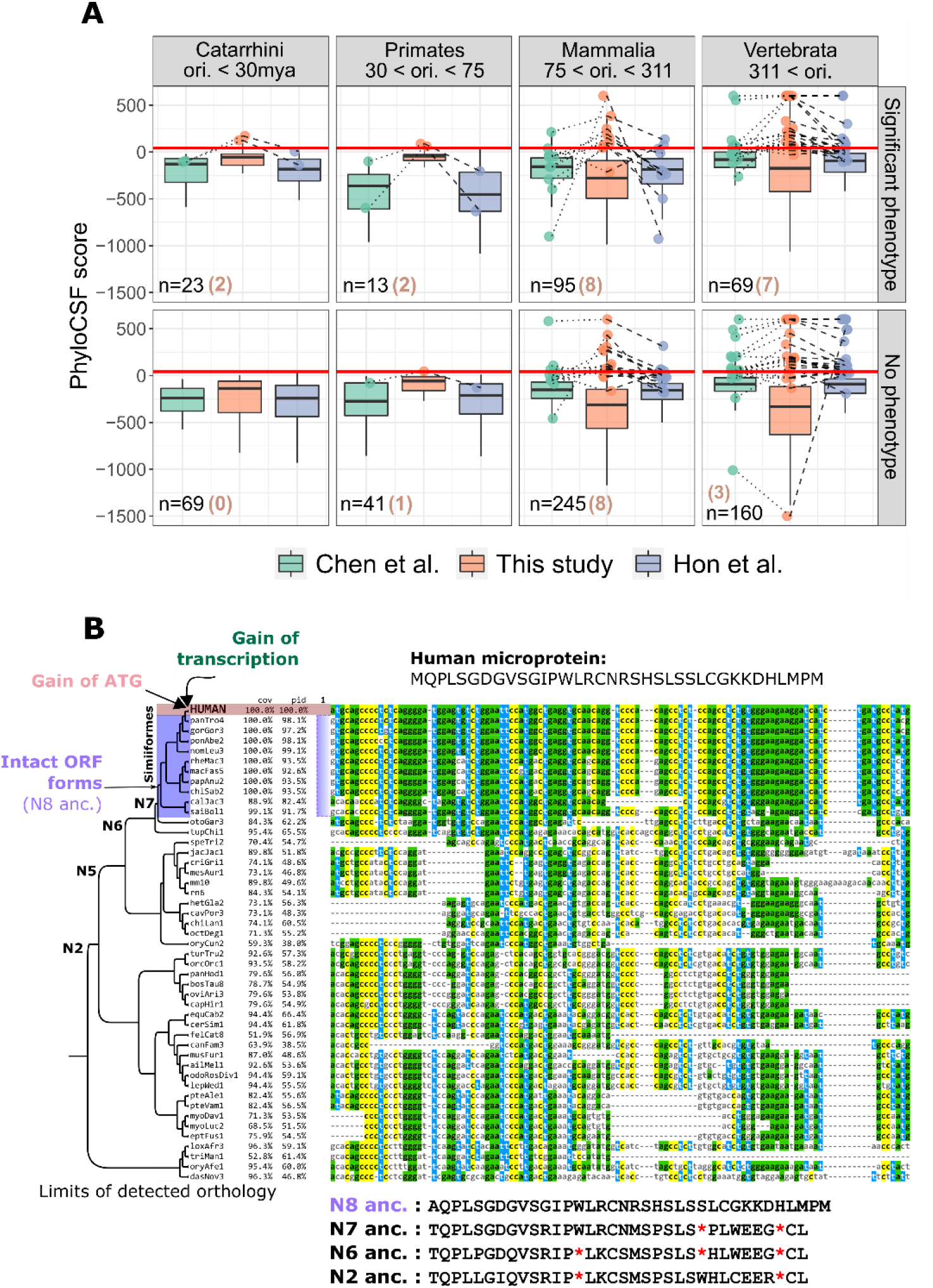
**A:** Boxplots show distributions of PhyloCSF scores as calculated in this study, by Hon et al. and by Chen et al. for all 715 microproteins (both de novo and undetermined origin) with and without significant phenotypes, (see Methods). Boxplot outliers are not shown. Maximum and minimum values have been set to 600 and −1,500 respectively to improve visualization. Points show microproteins that are predicted as coding (score ≥ 41, red horizontal line) by at least one study. Lines connect points corresponding to the same ORF. Microproteins are grouped in four broad classes of putative origin age. Numbers in parentheses are coding ORFs uniquely identified by our approach. **B:** Phylogenetic tree and multiple sequence alignment of ORF CATP00001771233.1 and its orthologous region in all species where it could be identified. Sequence names correspond to species assembly versions. The translated sequences of human ancestors in the +1 reading frame are shown. N5 ancestor is predicted to be identical to N6. Alignment visualized with Mview^50^, phylogenetic tree with FigTree.

Further evidence for the biological significance of a gene can come from an observed association with disease. The disruption of functionally relevant peptides could potentially have pathogenic consequences and even be of clinical importance. To identify such cases, we surveyed all known SNPs annotated as pathogenic or likely-pathogenic in dbSNP^46^, found within the boundaries of our ORFs’ exons.

We identified three such SNPs (summarized in Supp Table 3). ORF CATP00000063293.1 (upstream, de novo emerged with a putative origin at Simiiformes) contains one pathogenic SNP (rs1555735545, Single Nucleotide Variant), associated with Limb-girdle muscular dystrophy. The SNP is annotated as an Intron/5’ UTR variant, but does in fact also change the start codon of the encoded protein sequence (ATG -> ATA). Consistent with a possible functional role, this ORF has strong PhyloCSF signal, but only when calculated using our ORF-origin-aware approach (88.6 vs. −99, Chen and −205, Hon); and a very high phenotypic score (69, 87^th^ percentile of all ORFs screened by Chen et al.) in K562 cells.

The second pathogenic SNP is found on “new” ORF CATP00000005301.1 (SNV, G>A in the forward strand, rs1238109100). It is tagged as “Likely Pathogenic” related to Retinitis pigmentosa^47^. This protein is longer (178aa), predicted to be entirely disordered and it too has very high phenotypic score in K562 cells (47.2). It originated in Boreotheria and the lncRNA gene is most associated with melanocytes (source: Hon et al.). The mutation would change the 155^th^ amino acid from Histidine to Tyrosine. Once again, our ORF-origin-aware way of calculating PhyloCSF produces a strong coding prediction (1,959.299), whereas previous estimates had negative scores (−901 Chen, −927 Hon). The strength of this score was surprising, since the microprotein has a more ancient origin than the one described in the previous paragraph. We thus ran PhyloCSF again, on a normal, codon unaware alignment. As expected, the score was strongly negative (−3,062), stressing the importance of reading frame consideration in this type of alignment. CPAT applied on the ORF sequence only also predicts this ORF as coding, with a hexamer log-likelihood score of 0.19 (positive values indicate a coding sequence, negative values indicate a noncoding sequence) and a coding probability of 0.85.

The third SNP overlaps ORF CATP00000363722.1 (rs1560929898) and is a single nucleotide deletion that would cause a frameshift after the 16^th^ amino acid. The mutation is associated with Alazami syndrome^48^ which is in line with the ontology association of this lncRNA (embryonic stem cell related according to Hon et al.). Curiously, no significant phenotype or coding signatures were detected for this ORF. Yet we predict an ancient origin (Euteleostomi), and subcellular localization to the mitochondria. Note that, contrary to the first case, the effects of the second and third SNP could also be due to change in proteins produced by overlapping genes CDH3 and LARP7 (all potential consequences can be found in Supp. Table 3). Overall, these three cases provide excellent candidates for further exploration of the clinical significance of novel microproteins.

### A novel ORF epitomizes the “out-of-the-box” functional potential of de novo gene birth

We sought a clear-cut case to exemplify the capacity of de novo gene birth to readily produce a functional protein product. CATP00001771233.1 is a 108nt ORF, found on the intergenic lncRNA RP3-527G5.1 (ENSG00000231811.2; Chen et al. peptide RP3-527G5.1_4347298_36aa), which according to Hon et al. is transcribed through the action of an enhancer (e-lncRNA). The lncRNA gene is strictly human-specific and does not overlap other genes in any strand, with the exception of an intronic region of lncRNA gene ENSG00000285424 (there are however no overlapping exons, see Supp. Figure 3). This is confirmed by the data of Hon et al. (no orthologous transcription detected in any tissue in mouse, dog, rat, or chicken), by RNAcentral (taxonomy results show the transcript as only present in human) and by ENSEMBL (gene has 0 orthologs and is described as novel).

Based on its reported expression pattern (Hon et al.), the gene is strongly associated with heart tissue (ontology with strongest association is cardiac chamber, followed by cardiac valve, cardiac atrium, melanocyte, atrioventricular valve, and pigment cell, Supp. Figure 3). A very similar expression pattern is found in GTEx for this gene (most expressed in heart, by a large margin, Supp. Figure 3). The gene is also strongly differentially expressed during melanocytic induction, as well as two other experimental series.

The identification of the orthologous genomic region but lacking the ORF in species as evolutionarily distant as the armadillo; the results of the ASR; combined with the fact that the protein has no significant matches in any vertebrate proteome (or anywhere else in NCBI’s nr database), strongly suggests that this ORF emerged de novo (Figure 2B). Our conservative prediction is that the ORF formed at the ancestor of Simiiformes (using a 0.5 length ratio cut-off places the origin slightly earlier, at the ancestor of Primates, N6 in Figure 2B). The ATG start codon formed in the human branch, while all other primate species have a GTG codon at that position (Figure 2B). No tool predicts coding potential for this ORF or transcript (PhyloCSF Chen et al. score: −327.4246, PhyloCSF Hon et al. score: −318.1374, PhyloCSF our score: −54.3, max CPAT score of transcript: 0.072, CPAT hexamer score for ORF: 0.1, RNAcode p-value: 1, sORFs.org FLOSS score: −1). Nonetheless, the ORF is translated with high-confidence (ORF-RATER score of 0.85 in iPSC) and has a strong fitness effect in K562 cells (phenotype score: 61.2, 85^th^ percentile). Thus, we are confident both that this is a genuine gene, and that it emerged de novo uniquely to humans.

Given the association with heart, we also sought to confirm expression of this microprotein within a recent dataset of the heart translatome^49^. In this dataset too, the transcript is found to be human-specific as it is absent in rat and mouse. In human, both transcript and ORF show heart-specific expression (RNA only expressed and ORF only translated in iPS-derived cardiomyocytes). Furthermore, our analysis suggests that the ORF encodes an entirely disordered peptide that has predicted extracellular or nuclear localization. Overall, this example demonstrates that a newly emerged, human-specific ORF can be readily functional, under a highly specific expression program.

### Properties of young and ancient microproteins

In many organisms, it has been shown that evolutionarily novel genes have distinct sequence properties such as low expression and short length^39^. Although our dataset is biased since it only includes unannotated and thus shorter ORFs, we investigated potential differences in various ORF, transcript and protein properties across four different phylogenetic groups of origin of microproteins (Figure 3).

**Figure 3.**
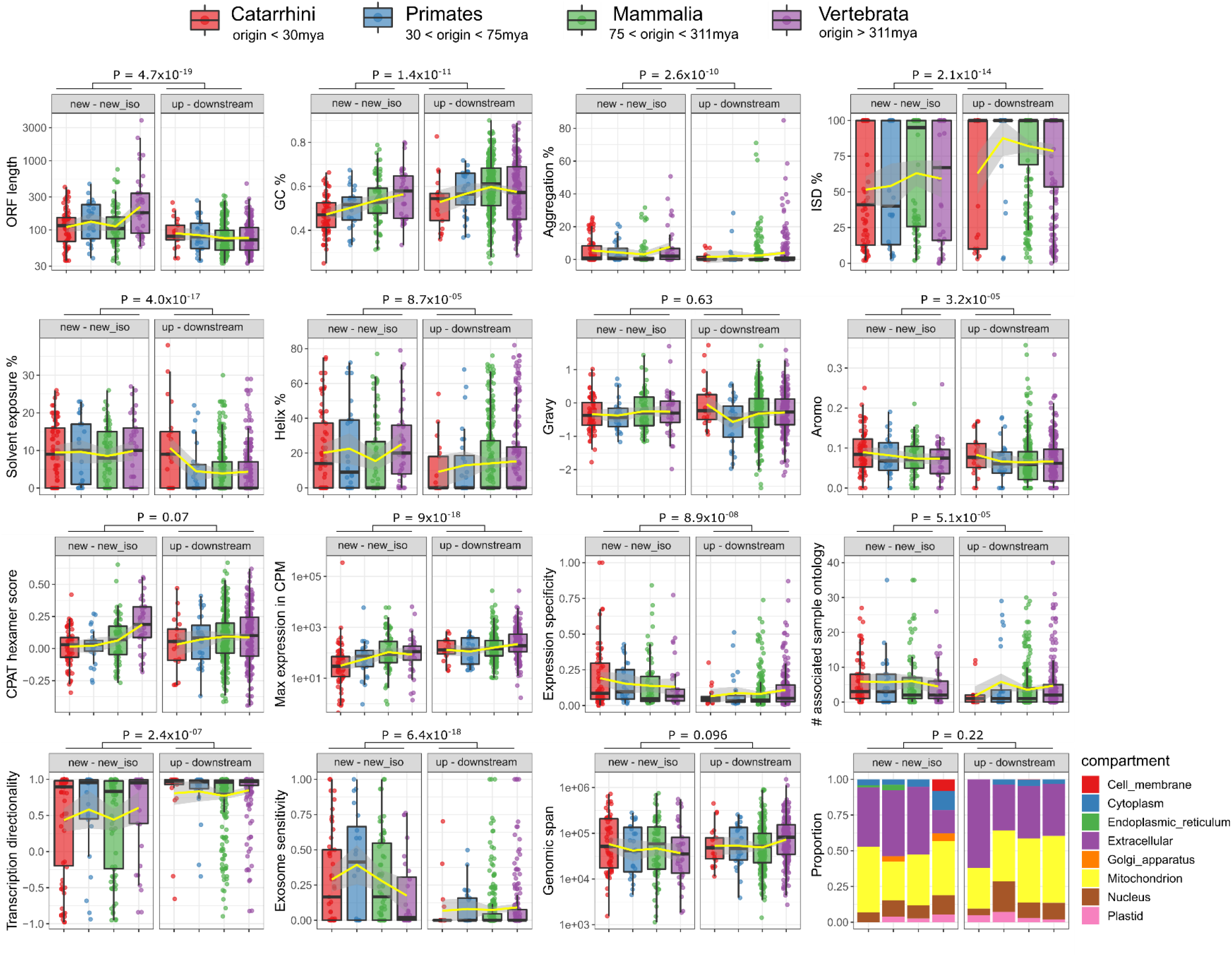
Distributions of various ORF, transcript and protein properties for all 715 microproteins, in four broad groups of putative origin age. Wilcoxon test P-values are shown for comparisons of all “new – new_iso” ORFs (n=211) to all “up - downstream” ones (n=504), except for subcellular localization (bottom right, X^2^ test). Yellow line shows LOESS fit and transparent frame around the line shows margin of error estimates.

The most significant difference was observed for GC content between ORFs in the Catarrhini and Vertebrata groups, especially for “new – new_iso” ORFs (avg. GC% 0.47 vs. 0.56, Wilcoxon’s test P-value=0.00012). Indeed, there is a correlation between GC content and time since putative origin (Spearman’s Rho 0.31, P-value=3.4×10^−6^). The difference is not statistically significant for the “up - downstream” class (avg. GC% 0.53 vs. 0.57, Wilcoxon’s test P-value=0.13), and there is no correlation of GC% to time since origination (P-value=0.76). A similar trend (young ORFs having lower GC% than older ORFs) was observed by Dowling et al.^38^, albeit between slightly different phylogenetic groups.

An equally significant difference was found when comparing the Hexamer score (nucleotide hexamer usage bias between coding and noncoding sequences) calculated by CPAT for ORFs. Again, the difference is only found for “new - new_iso” ORFs (Avg. Hexamer score 0.016 vs. 0.19, Wilcoxon’s test P-value = 5.3×10^−6^) and absent in the “up - downstream” class (P-value = 0.26). This could reflect a tendency in “new” ORFs to become more “gene-like” and more optimized with time. Such a tendency could be absent in up - downstream ORFs since they are expected to be enriched in sequence-independent function.

Comparing all “new – new_iso” ORFs to all “up - downstream” ones reveals differences in most features we explored. Somewhat expectedly, “new – new_iso” ORFs are longer, GC poorer (albeit that these two properties probably correlate given the fact that start and stop codons are AT rich^39^), the transcripts they are found on are less expressed and with higher tissue specificity, they have more associated sample ontologies, have lower transcriptional directionality and are more exosome sensitive. They encode microproteins with higher aggregation propensity, less intrinsic disorder, higher solvent accessibility, higher helix content and more aromatic residues (Figure 3). Interestingly, only 15 out of the 715 ORFs were found to encode a TM domain showing no timing of origin or significant phenotype bias (X^2^ test, both P-values > 0.14). This contrasts with recent findings from budding yeast where the propensity to form transmembrane domains is prevalent among young de novo genes^26^. No significant difference was observed in predicted subcellular localization between the various classes.

## Discussion

Ribosome Profiling has enabled the accurate identification of translated sORFs. Coupled with experimental evidence for the phenotypic effects of the encoded microproteins, this presents an excellent opportunity to study the evolutionary origin of these elements without being limited by the use of conservation as a proxy for function. Here we explored the set of seemingly functional microproteins and uncovered strong evidence of unequivocal cases of recent de novo origination.

Our conservative estimate is that at least 6 such biologically significant human-specific microproteins exist, with additional ones having a slightly older, but recent, origin, since the ancestor of all primates. Additional, more targeted experimental work is now needed to conclusively demonstrate both the functional role of these elements in human, but also the absence thereof in other species. Such a confirmation, if it arrives, would be especially consequential for how we annotate functional coding regions in the future. We have no theoretical reason to doubt that a functional microprotein can exist in almost total absence of detectable evolutionary constraints due to its recent origin. This scenario creates a need for models that take such conditions into account. Given how rich the human translatome, transcriptome and ORFeome are, meeting this need could prove challenging.

An important question is why, out of all the translated novel ORFs, some rapidly acquire biological function while others do not. This is essential to identify the biologically relevant sORFs out of the potentially thousands that could be translated^5^. Will future advances reveal a protogene-model-like reality^25^, where, out of a wide pool of candidates a few functional ones evolve stochastically? Or could natural selection have acted already, to enrich translation of ORFs already more or less primed for functionality, for example by eliminating those likely to be toxic^51^? Are these novel microproteins always recruited in a functionally specific manner, or do they initially have more generalized roles? Plausibly, tissue-specific expression of novel peptides might initially carry a smaller risk of overall deleterious effects, thereby potentially “shielding” them from the action of purifying selection. It will also be interesting to understand if, and why, some microproteins evolve in terms of their length, sequence properties and functional role, while others remain unchanged, resembling frozen accidents^52^. Here, we found only limited differences between young and ancient, but additional efforts and larger datasets will be needed to explore this in the future.

A related, widely discussed question in the field of de novo gene origination is whether the genomic loci out of which novel ORFs emerge have particular characteristics. A high GC-content would favor the formation of ORFs, as stop codons are GC-poor, however we did not find that young ORFs are more GC-rich than ancient ones, rather the opposite. This could be partially explained by a selective pressure for increased GC-content (and thus increased expression and/or nuclear export) acting on intronless genes^53^, which are overrepresented among the ORFs studied here (459/715 are single exon). Another possibility is that novel ORFs may emerge out of pseudogenic loci, where longer ORFs once existed but are now defunct. Curiously, the region of CATP00001771233.1, the exemplar ORF above, could correspond to something similar. While the first frame of the ORF produces a negative PhyloCSF score, the second frame produces a positive score of 48. We did a search for ORFs on the full sequence of the transcript downloaded from ENSEMBL and found that there are multiple longer overlapping ORFs, including an 85aa one in the reverse strand. None of them are found as translated by Chen et al. and 3 additional ones (apart from CATP00001771233.1) are found translated by van Heesch et al., but these do not correspond to the longer existing ones. More detailed investigation of the genomic region is needed to fully appreciate the meaning of this observation.

The true proportion of young de novo ORFs with phenotypic consequences could be higher. Our study is limited in using data from only one series of knockout experiments, but a wider, more comprehensive analysis could bring a more accurate view into focus. This would also increase confidence in the phenotypic effects that are detected, decrease the number of false positives, and allow to more safely build on top of these findings. It is plausible that ancestral sequence reconstruction might also be a source of false positives, in which case some of the ORFs we identified as de novo emerged could correspond to ancient ones which, through a combination of factors such as sequence divergence, deletions or chromosomal rearrangements, appear recent. Application of multiple ASR methodologies and thorough assessment of reconstruction uncertainty could in the future alleviate such problems and increase accuracy.

## Materials and Methods

### Data collection

Our dataset included ORFs that were identified as translated with high confidence by Chen et al.^40^ based on analysis performed by the ORF-RATER program^54^ (ORF-RATER score >= 0.8). Following the classification of Chen et al., we restricted our analysis to those ORFs located on either previously annotated non-coding transcripts (“new”), upstream of coding ORFs on coding transcripts (“upstream“), downstream of coding ORFs on coding transcripts (“downstream”) or on transcripts lacking coding ORFs but which belong to a transcript family with a member annotated as coding (“new_iso”). We also required that ORFs be present in the comprehensive catalogue established by Hon et al.^41^ (FANTOM-CAT dataset). ORFs from the two studies were matched based on identical chromosomal coordinates, 100% sequence identity and identical length. Our final dataset consisted of 715 ORFs, located on 527 unique transcripts. Note that some of these ORFs overlap with others. Human genome version *hg19* coordinates were converted to *hg38* using the *liftover* tool in UCSC Genome Browser.

Various types of data were collected and generated for each ORF and its encoded protein. We considered whether the ORF was found to have significant fitness effects according to Chen et al.’s high-throughput CRISPR-Cas knockout screens in iPSC and K562 chronic myeloid leukemia cells. Phenotypic scores and classification (significant/not significant) were collected from the data of Chen et al. for each ORF in the two cell lines. Orthologous transcription, various coding signatures for ORFs and transcripts, expression data, cell type association, trait association, transcription properties for each transcript were obtained from the supplementary data and the raw data depository of Hon et al.^41^ Protein secondary structure was predicted by RaptorX^55^ using default parameters, transmembrane domains were predicted with Phobius^56^, disordered regions were predicted with IUPRED^57^, subcellular localization was predicted with DeepLoc^58^ and percentage of aromatic and hydrophobic amino acids were calculated with codonw^59^. CPAT^44^ was applied on the sequences of the ORFs to calculate the Hexamer and coding probability scores.

### Identification of orthologous genomic regions and orthologous transcription

For each of the 715 human ORFs, we identified its orthologous region in 99 vertebrate genomes based on the UCSC Genome Browser 100-way, whole-genome alignments. The exact orthologous genomic regions corresponding to each exon of the human ORF were extracted using custom Python scripts. The regions corresponding to the different exons were then stitched together. For all ORFs, the orthologous region could be identified in a minimum of 4 other genomes. A multiple sequence alignment of each ORF together with its orthologous sequences was then performed using MAFFT^60^.

We inferred orthologous transcription by two means. First, we downloaded the NCBI RefSeq annotation GTF files for 92/99 vertebrate species for which it was available. Before it was possible to detect whether orthologous regions of ORFs were transcribed however, it was necessary to convert genomic coordinates from the assembly versions used in the 100-way alignments, to those used in RefSeq. To do this, we performed BLASTn^61^ searches of the orthologous regions to their corresponding genome RefSeq assemblies using a cut-off of 97% identity. We were thus able to define the coordinates of each exon in the RefSeq version of the assemblies. We then verified that all updated coordinates produced in this manner were indeed as close as expected to the previous ones (i.e. we confirmed that no irrelevant matches were retrieved). We then checked, in each species, whether at least one exon of each orthologous region overlapped with an annotated transcript. An 80% overlapping cut-off was used. This gave us an initial pattern of presence and absence of transcription across the 92 species. Furthermore, we performed additional BLASTn searches of each human ORF to the entire vertebrate RefSeq transcript sequence database (downloaded March of 2021 from NCBI’s ftp website, https://ftp.ncbi.nlm.nih.gov/refseq/release/vertebrate_mammalian/plus https://ftp.ncbi.nlm.nih.gov/refseq/release/vertebrate_other/) and considered the region as transcribed if it matched at the sequence level (regardless of genomic position) any transcript in a given species, with at least 60% query coverage and at least 0.001 E-value. Note that only very rarely a presence was inferred in this manner that was not also retrieved based on annotation overlap. Based on each ORF’s phyletic pattern of transcription presence and absence on 92 vertebrate species, we were able to define the transcriptional origin by taking the most recent common ancestor of all the species for which a presence was inferred (Dollo parsimony). To do this, we used the 100-way phylogenetic tree from the UCSC genome browser. Phylogenetic nodes were then matched to their official taxonomic names through TimeTree.org^62^.

### Ancestral Sequence Reconstruction and inference of presence of ancestral ORFs

For each ORF, we first pruned the UCSC 100-way phylogenetic tree using the *gotree* tool^63^ to keep only leaves corresponding to species present in the alignment of orthologous regions. A phylogenetic tree following the pruned tree’s species topology was then constructed from the multiple alignment of each ORF and its orthologous sequences, using RAxML^64^ (*raxml-ng --evaluate --msa ORF_alignment.fa --model GTR+G --tree pruned_ORF_tree.nwk*). Then, each multiple alignment and its corresponding tree were given as input to FastML^65^, to reconstruct the various ancestral sequences. The JC substitution matrix was used, and the ML method was used for reconstruction of indels. The marginal ancestral reconstructions were then parsed.

We examined the reconstructed sequences of the human ancestors. The origin of the human ORF was defined as the most ancient human ancestor in which at least 70% of the reconstructed ancestral sequence was an intact ORF (length of ancestral ORF/length of full ancestral sequence), i.e. any premature stop codons did not disrupt more than 30% of the length of the sequence (a 50% and an 80% cut-off was also used, see main text for details). The reading frame used was always on the forward strand, starting from the first position of the reconstructed sequence. If the ancestral sequence was longer than the human one, the length of the human sequence was used as the denominator of the ratio (length of ancestral ORF/length of human ORF). If the length of the reconstructed sequence was less than half the length of the human ORF, the ancestor was not taken into account, effectively considered as intact. Ancestral sequences were counted as intact ORFs regardless of whether an ATG start codon was present or not. We distinguished cases for which at least one disrupted (not intact) ancestor could be identified on a more ancient node than the one of predicted origin. For these cases, we were thus able to provide positive evidence of de novo formation: a disrupted ancestor that preceded the most ancient intact one. To be maximally conservative, we also conducted protein level similarity searches of all candidate ORFs, using BLASTp, against the annotated proteomes of all “outgroup” species, i.e. those diverged prior to the predicted node of origin of the ORF (proteomes downloaded from NCBI’s RefSeq). Matches were deemed as significant if they had < 10^−5^ E-value, 40% identity and 50% query coverage. Based on these matches, we reassigned the node of origin to the most recent ancestor of the expanded set of species when necessary and removed de novo origin status. This was applied to 17 cases.

To detect possible inconsistencies with de novo origination, we performed a search for similarity to already annotated human protein sequences (Homo_sapiens.GRCh38.pep.all.fa file downloaded from ENSEMBL) using BLASTp with an E-value cut-off of 10^−5^, 50% identity and 50% query coverage, providing as query our de novo originated ORFs. We recovered two matches. One of them was the protein itself (CATP00000191117.1 -> ENSP00000493702.1, 100% identity). We confirmed from ENSEMBL that the annotated gene (ENSG00000170846, which has only one protein associated) had the same predicted origin (Eutheria, from the Gene gain/loss tab) as the one we calculated, for both ORF and transcript. According to ENSEMBL, the gene also has two paralogues, which both originated at the root of Eutheria. The second match came from ORF CATP00001059838.1. This ORF again matches part of an ORF of its own gene (ENSG00000267360), but at 76% identity. The origin of the gene is more ancient (Boreoeutheria) than our predicted origin of the ORF (Simiiformes), but this is expected since this is an upstream ORF, and not the main coding ORF of the transcript. Overall, this search revealed no inconsistencies linked to human paralogues of our candidates.

### Functional signatures and statistics

We extracted all SNPs from dbSNP^46^ within the coordinates of each of our ORFs that were not annotated as benign, using the following command, for each exon:

> *esearch -db snp -query CHR_NO AND (START_COORD:STOP_COORD) NOT “benign”[Clinical Significance])” | efetch -format json*

Detailed information for each SNP was then retrieved from the SNP’s page at dbSNP and ClinVar.

To calculate PhyloCSF^42^ scores, we placed the human ORF sequence and orthologous sequences in species descending from the phylogenetic node of origin of the ORF in a FASTA file. We took the origin of the ORF and not the putative origin of the microprotein to minimize cases of origin age underestimation due to incomplete transcript annotation, as mentioned in the main text. We then generated a codon-aware nucleotide alignment with the TranslatorX^66^ tool, keeping the reading frame unchanged. PhyloCSF scores were then calculated based on these alignments, using the human sequence as reference and the −removeRefGaps option, searching only in the first reading frame and employing the “vertebrates100” model. PhyloCSF was applied by Chen et al. on alignments including sequences from 10 mammals spanning the Euarchontoglires, and by Hon et al. on alignments including 27 mammalian species. All statistics were done in R version 3.6.2. Plots were generated using ggplot2^67^.

## Data availability

Raw data and scripts that generate the figures are available **(*will be once the work is published*)** at https://github.com/Nikos22/humandenovo

## Acknowledgments

We are grateful to Jin Chen for providing us with data and methodological details from ^40^. We also thank Laurence Hurst and Anthony Redmond for valuable feedback on the manuscript. This work was supported by funding from the European Research Council, grant agreement 771419. This research is co-financed by Greece and the European Union (European Social Fund-ESF) through the Operational Programme «Human Resources Development, Education and Lifelong Learning» in the context of the project “Reinforcement of Postdoctoral Researchers - 2nd Cycle” (MIS-5033021), implemented by the State Scholarships Foundation (IKY).

## Author information

### Contributions

AMcL and NV conceived the study. NV performed the analyses. NV and KD analysed the data. AMcL and NV wrote the paper. All authors read, finalized and approved the final manuscript.

### Corresponding authors

Correspondence to Aoife McLysaght and Nikolaos Vakirlis

## Supplementary Figures

**Supplementary Figure 1.**
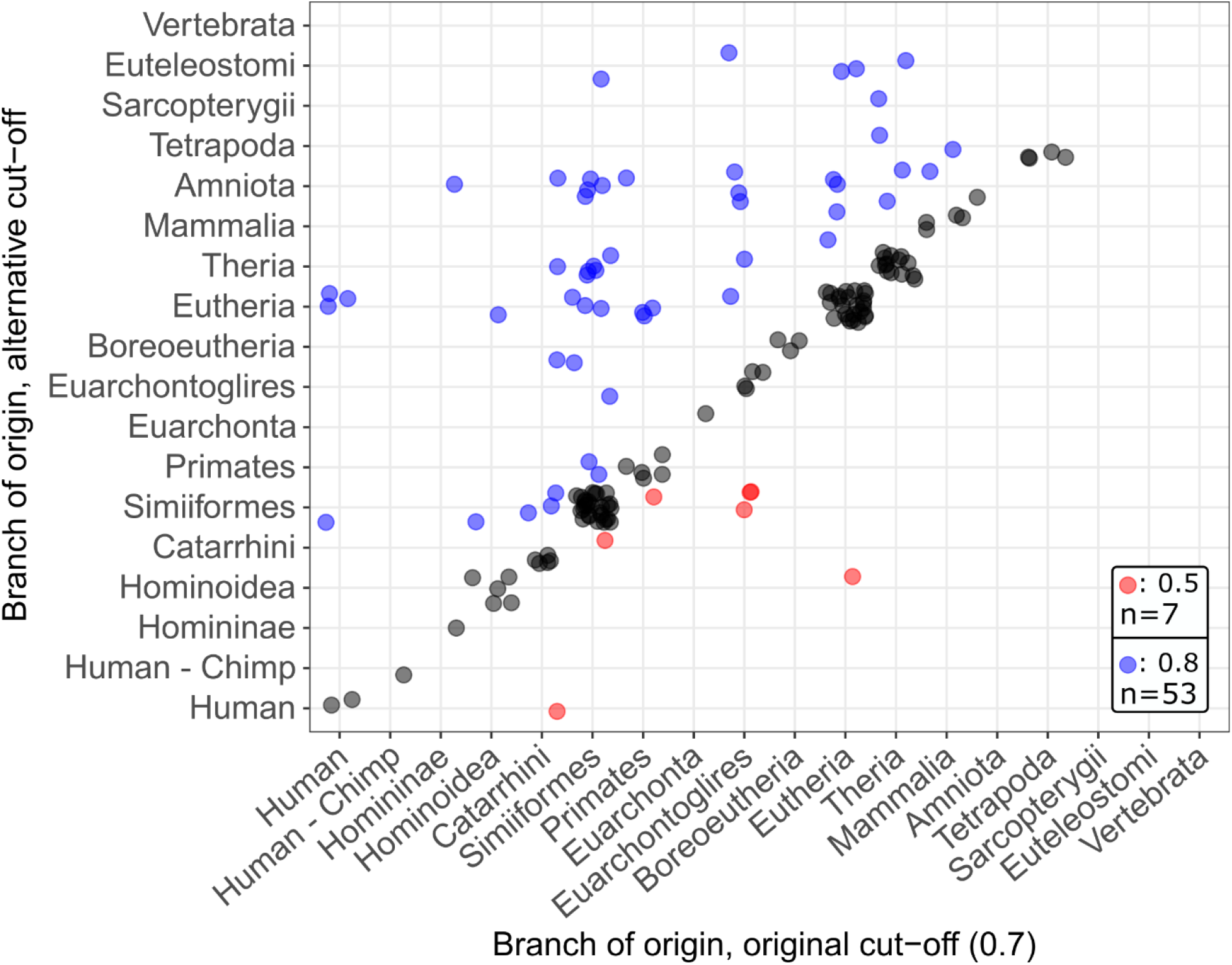
Effects of applying two alternative intact length proportion cut-offs, one stricter (red points) and one more relaxed (blue points) on predicted branch of ORF origin for 155 de novo originated ORFs.

**Supplementary Figure 2.**
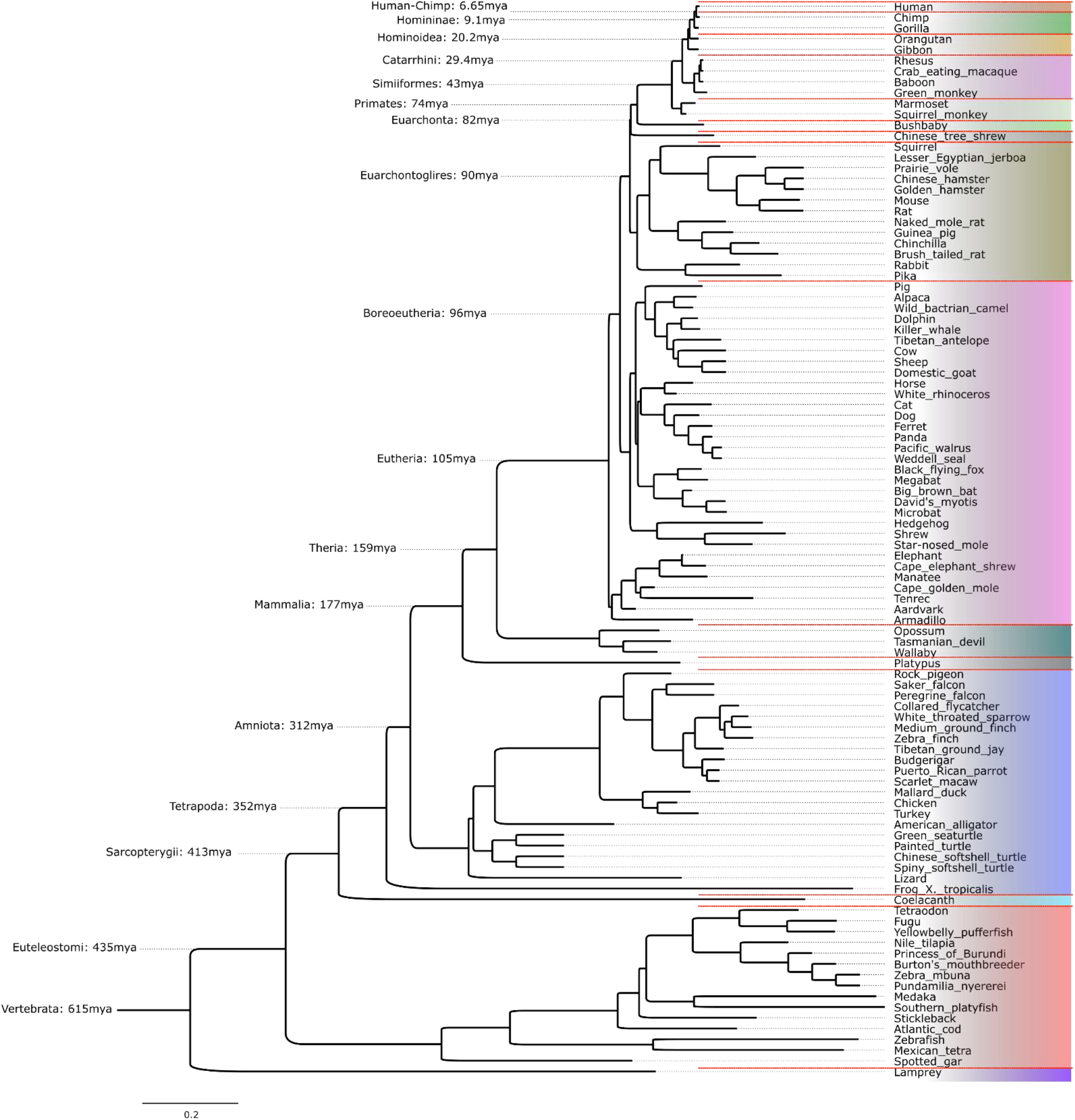
UCSC Genome Browser 100-way phylogenetic tree with common species names annotated with the human ancestral branches and their ages, as estimated by TimeTree.

**Supplementary Figure 3.**
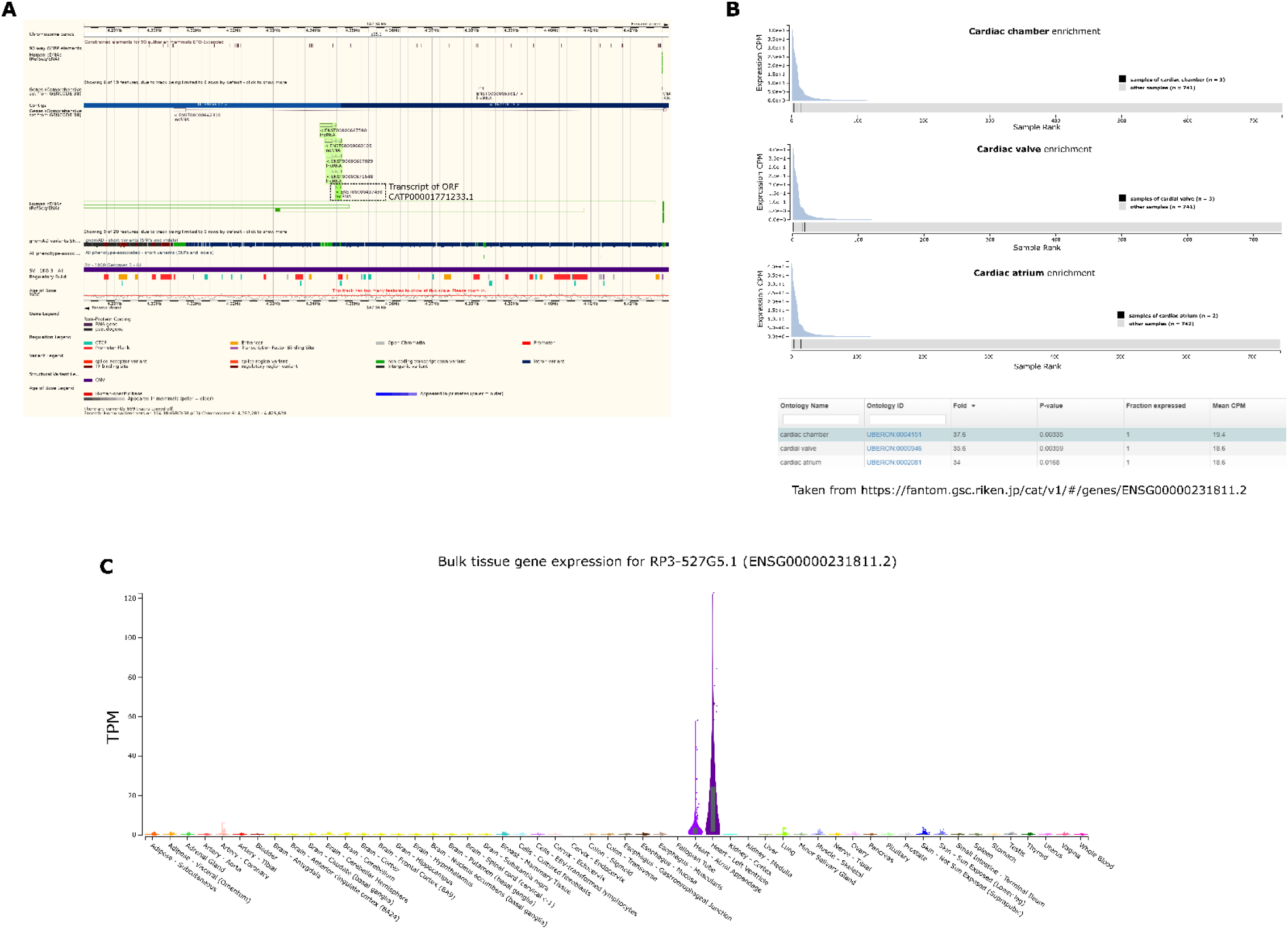
A: Genome browser view from ENSEMBL of the region of gene ENSG00000231811.2. The transcripts of the gene are highlighted in green. B: Sample ontology enrichment (three highest ranking ones) detected by Hon et al. for gene ENSG00000231811.2. Plots show the ranks of the samples for each ontology, table shows the relevant information. Taken from https://fantom.gsc.riken.jp/cat/v1/#/genes/ENSG00000231811.2 C: Expression plot taken from GTeX taken from https://gtexportal.org/home/gene/RP3-527G5.1#geneExpression.

## Supplementary Tables

**Supplementary Table 1.**

Species names and assemblies of vertebrate genomes used in this study.

**Supplementary Table 2.**

Data for the 715 ORFs analysed in this study.

**Supplementary Table 3.**

Information on three SNPs overlapping ORFs analysed in this study.

